# Accurate Mutation Effect Prediction using RoseTTAFold

**DOI:** 10.1101/2022.11.04.515218

**Authors:** Sanaa Mansoor, Minkyung Baek, David Juergens, Joseph L. Watson, David Baker

## Abstract

Predicting the effects of mutations on protein function is an outstanding challenge. Here we assess the performance of the deep learning based RoseTTAFold structure prediction and design method for unsupervised mutation effect prediction. Using RoseTTAFold in inference mode, without any additional training, we obtain state of the art accuracy on predicting mutation effects for a set of diverse protein families. Thus, although the architecture of RoseTTAFold was developed to address the protein structure prediction problem, during model training RoseTTAFold acquired an understanding of the mutational landscapes of proteins comparable to that of large recently developed language models. The ability to reason over structure as well as sequence could enable even more precise mutation effect predictions following supervised training.

## Main Text

Accurate and unsupervised prediction of single mutation effects using sequence information alone would help relate observed sequence polymorphisms to human disease [1, 2] and contribute to the design of proteins with higher functional activities. Deep learning methods have recently shown considerable promise for mutation effect prediction. DeepSequence [3], a probabilistic model for sequence families, obtained excellent performance in mutation effect prediction using latent variables for capturing higher-order interactions between residues in proteins through training on multiple sequence alignments (MSAs) for the target protein of interest. Large protein language models trained on multiple sequence alignments (MSA Transformer) [4] or single sequences [5] also performed very well at mutation effect prediction, and have the advantage over DeepSequence of not requiring specific training on the protein of interest. RoseTTAFold was originally developed for protein structure prediction [6], but during training we included a masked token recovery task, and a recently developed version, RoseTTAFold Joint (RF_joint_) was further trained to solve ‘inpainting’ problems in which substantial portions of both sequence and structure are rebuilt [7]. To assess RF_joint_’s understanding of protein sequence-structure relationships, we set out to investigate whether it could predict experimental mutational data from published deep mutational scanning (DMS) sets [8] with no further training (using a zero-shot approach). We compared RoseTTAFold performance on this task to that of the state of the art MSA Transformer; both are MSA based methods requiring no further training.

RF_joint_ was evaluated on a set of 38 deep mutational scans curated by Riesselman et al. [3]. Each of the mutational scans recorded a different protein function with varying measurements. Each dataset was treated as a separate prediction task, and each variant was scored individually. For each target protein, we generated MSAs using iterative sequence search against the UniClust30 database as described in Baek *et al*. [6] and used it for both RF_joint_ and MSA Transformer predictions. For RF_joint_, the variants were scored by masking out the mutation site in the query sequence in the MSA and the MSA token recovery head was used to predict the distribution over the masked position. The predicted effect of the mutation was calculated as the log odds ratio of the mutant amino acid and the wild-type amino acid (Figure 1). The performance on each dataset was assessed based on the spearman correlation of the predictions to the observed experimental values.

**Figure 1.**
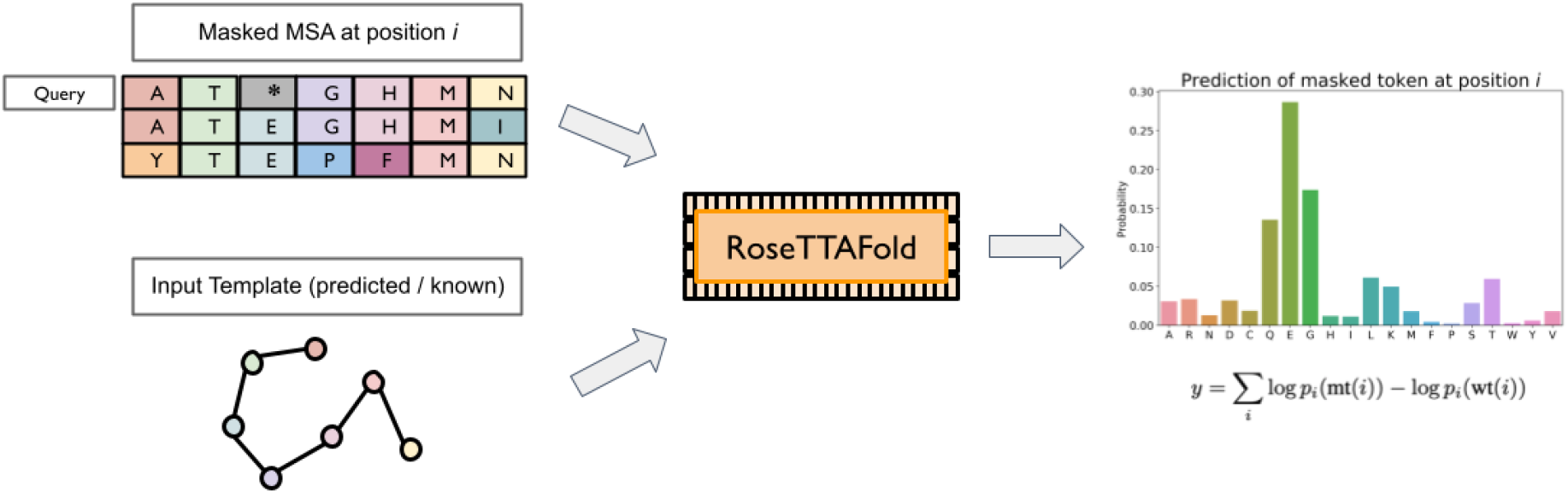
Overall pipeline for zero-shot prediction of mutation effect using RoseTTAFold. A MSA is generated and masked at the mutation position in the query sequence, and structural templates are fed into pre-trained RoseTTAFold. Using the masked token prediction head, the emitted probability distribution of the 20 amino acids over the mutation site is used to calculate the effect of a mutation as the log odds ratio of the wild-type and mutation amino acid.

We found that RF_joint_ predicts mutational effects considerably better than a baseline calculated as the log odds ratio of the frequency of the mutant amino acid and of the wild-type amino acid in the MSA (Figure 2). RF_joint_ also slightly outperformed MSA Transformer (Figure 2). RoseTTAFold has the advantage in principle over the purely sequence based models of also being able to utilize structural template information, but we did not observe a significant improvement with incorporation of template structure information (data not shown; this may be in part because RoseTTAFold generates 3D models from sequence with reasonable accuracy). We also found little dependency of prediction accuracy on MSA depth (Supplementary Figure 1).

**Figure 2.**
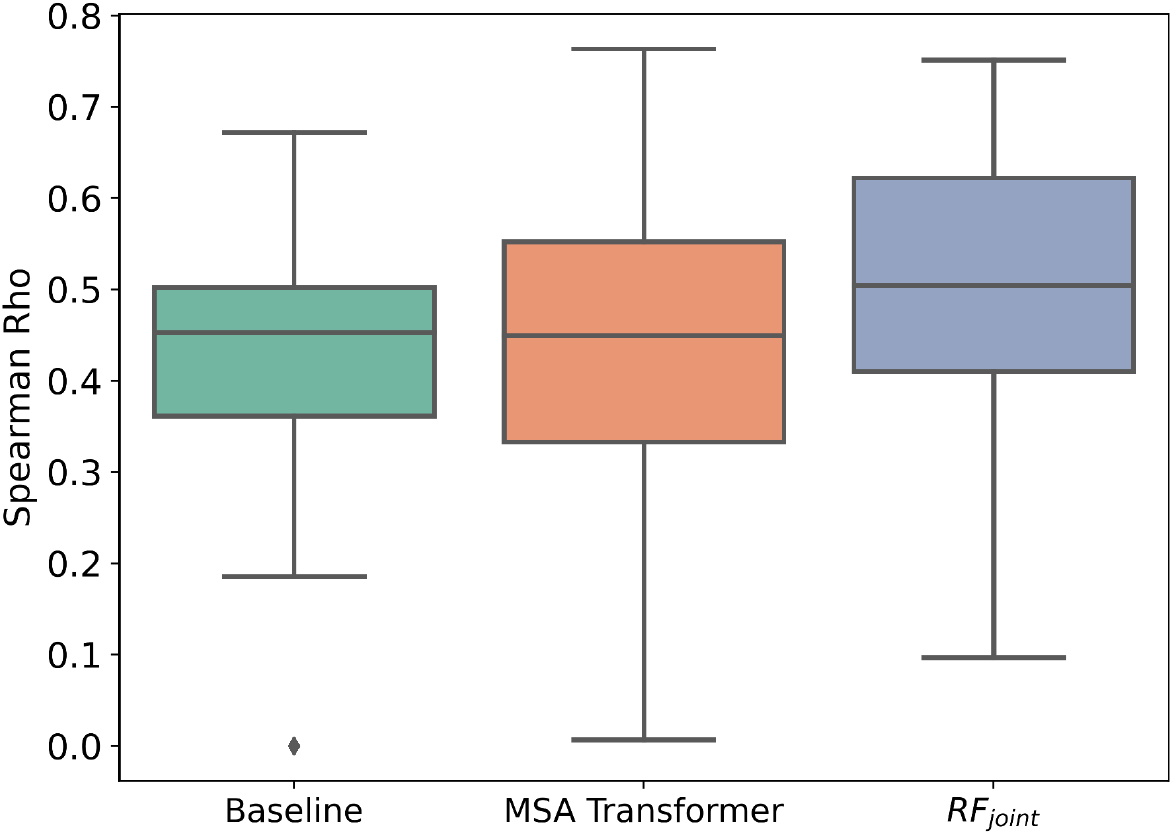
Boxplots of spearman rho correlations on deep mutation scanning datasets. Baseline refers to the non-ML MSA baseline. RF_joint_ refers to the model trained on a joint sequence and structure recovery task [7]. Box plots show the median (center line), interquartile range (hinges), and 1.5 times the interquartile range (whiskers); outliers are plotted as individual points.

## Conclusion

We find that the RoseTTAFold network, developed originally for structure prediction and then extended to protein design, is also able to predict the effect of single mutations with quite high accuracy. Just as large language models like the MSA Transformer provide general models of protein sequence, RoseTTAFold joint may be viewed as a general joint model of protein sequence and structure. With further more directed training, it should be possible to further improve performance by better utilizing protein structural information, which can be readily input into RoseTTAFold but not into pure sequence based models, and by fine-tuning specifically for the mutational effect prediction task. More generally, our results demonstrate that RoseTTAFold has quite a broad understanding of protein mutational landscapes, which should be very useful for protein design and other challenges involving inference over both sequence and structure.

## Materials and Methods

We used the published RF_joint_ model [7] in inference mode for the task of single mutation effect prediction. All weights of the model were frozen and no further training was done. Up to 256 sequences were considered from the input MSA of a target protein with an additional 1024 extra sequences passed into the model. All default parameters from RF_joint_ were used and the number of recycles was set to 1. RoseTTAFold [6] predicted structures for a target protein were used as structural templates for mutation effect prediction. Inference code for predicting the effect of single mutations through this pipeline is available here: https://github.com/RosettaCommons/RFDesign/tree/main/inpainting

## Acknowledgements and Disclosure of Funding

We would like to thank Justas Dauparas, Ivan Anishchanka, Doug Tischer, Hahnbeom Park, Sergey Ovchinnikov and Eric Horvitz for helpful comments and suggestions. This work was supported by Microsoft (S.M. M.B., D.B., and generous gifts of Azure computing time), Eric and Wendy Schmidt by recommendation of the Schmidt Futures (D.J.), the EMBO Postdoctoral Fellowship (ALTF 292-2022) (J.L.W.), the Audacious Project at the Institute for Protein Design (D.B.), a gift from Amgen (J.LW. and D.B.) and the Howard Hughes Medical Institute (D.B.).

---

**Supplementary Figure 1.**
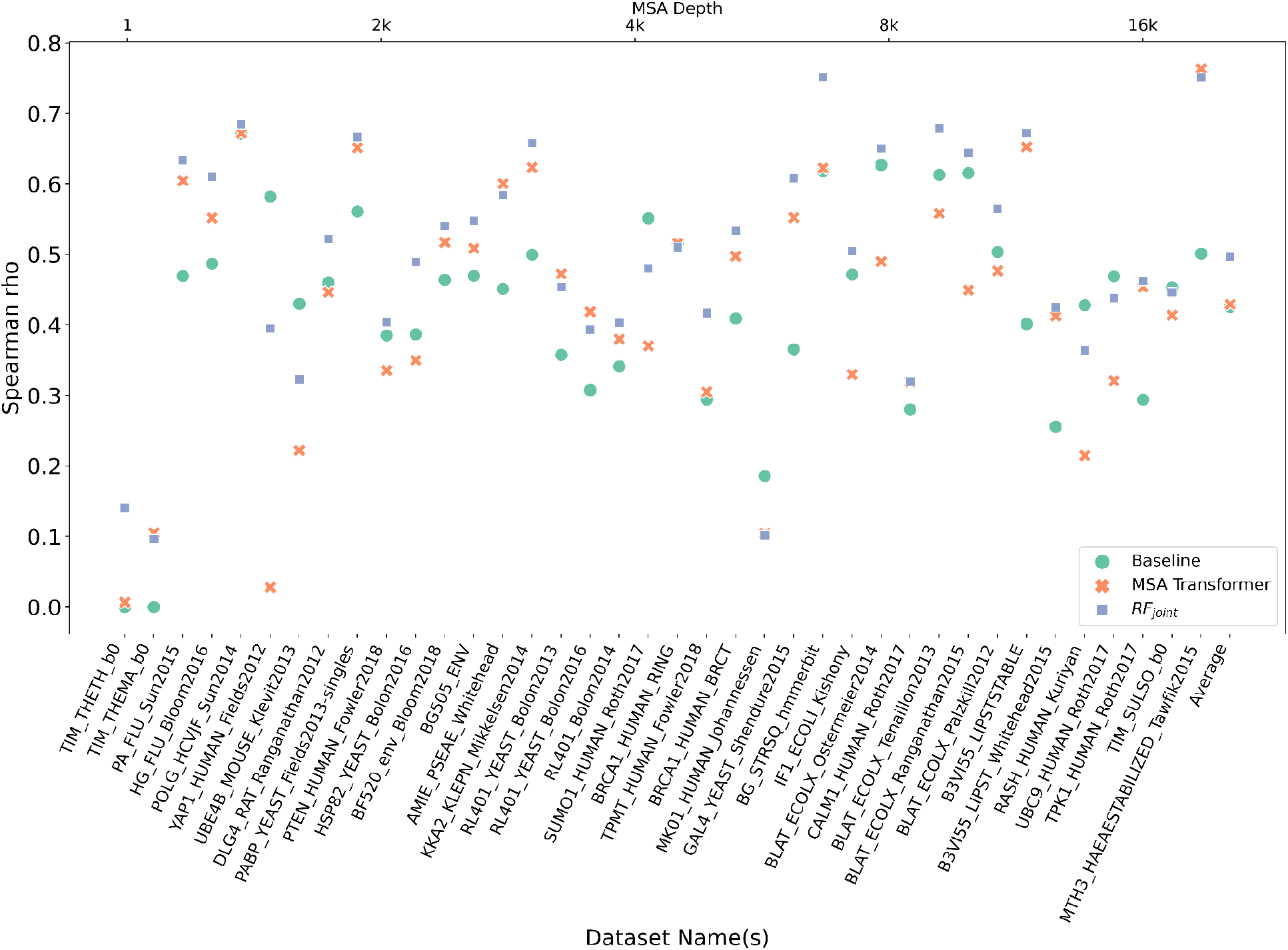
Spearman rho correlations for all deep mutational scanning datasets evaluated. Each point corresponds to a different protein. The points are arranged according to increasing MSA depth for RF_joint_ and MSA Transformer.

